# Melanocytes Exhibit Distinct Cell States Governed by A Gene Regulatory Network Under Stochastic Influence

**DOI:** 10.1101/2023.05.10.540111

**Authors:** Ayush Aggarwal, Ayesha Nasreen, Sarthak Sahoo, Mohammed Faruq, Rajesh Pandey, Mohit K Jolly, Abhyudai Singh, Rajesh S Gokhale, Vivek T Natarajan

## Abstract

Melanocytes serve as a protector of the skin against external stressors such as ultraviolet radiations. While melanocytes remain functional for nearly the entire lifespan of an individual, how they respond efficiently and survive such insults remain elusive. Here, we show the co-existence of multiple distinct states of normal human epidermal melanocytes (NHEMs). Using a progressive pigmentation model, we show that stochasticity in gene expression can lead to the co-existence of multiple melanocyte states. Using active enhancers and gene expression footprint, we identified state-specific transcription factors and constructed a gene regulatory network (GRN). The GRN couples pigmentation and cell cycle regulators and explains the co-existence and transitions of the melanocyte states. Finally, we show that NHEMs respond to external cues by altering the cell state dynamics. These results demonstrate that stochasticity in gene expression followed by epigenetic modification leads to the co-existence of multiple melanocyte states enabling an efficient response to environmental cues.

## Introduction

Heterogeneity within a seemingly homogeneous population is necessary to maintain the diversity required for the survival of the overall population. While this diversity is evident at the organism level, it was not observed/detectable at the cellular level until the last decade. Lately, with the advent of single-cell sequencing (scRNA seq) technologies, more and more evidence has revealed heterogeneity within a previously thought homogeneous pool of cells^1^. This has revolutionized the field of cancer research by unravelling previously unknown cell states of the transformed cancer cell^2–5^. Although heterogeneity was thought to be a consequence of malignant transformation, recent evidence points towards pre-existing diversity even in non-transformed cells. CD8+ T cells, microglia and fibroblast display distinct cell states under different conditions^6–8^. Variability in intrinsic cell states is proposed to be a generic feature, perhaps to enable faster and more efficient response to external cues^9^.

Human skin faces constant exposure to external cues such as UV, environmental pollutants, and microbial colonization. Epidermal keratinocytes deal with these insults by continuously shedding the outermost skin layer with constant replenishment^10^. However, the coping mechanisms of melanocytes present in the basal layer remain largely unexplored. Melanocytes are pigment-producing cell types derived from a transient multipotent population that emerge from the neural crest^11–13^. These cells occupy the basal epidermis and hair follicle in the human skin and impart their functional role of photoprotection by the production and transfer of melanin to keratinocytes and growing hair shaft, respectively^14–17^. In the hair follicle niche, melanocyte stem cell (McSc) transitions through multiple cell states, including a proliferative/transit amplifying (TA) state, to give rise to the mature pigmented state^18^. The TA state is a reversible state that dedifferentiates to the McSc state in the bulge region^19^.

Although NHEMs are not known to exhibit different states naturally, the transformation of melanocytes to a malignant state reveals the underlying plasticity. Melanoma exists in two broad states, melanocytic-like (Mel) and mesenchymal-like (Mes), that differ in their proliferation and invasion potential^20–25^. Recently, an intermediate state governed by a distinct chromatin landscape and a stable GRN was identified^26^. These multiple melanoma states are governed by complex multi-omics regulations, such as state-specific enhancers defined by H3K27ac and chromatin accessibility^27, 28^. These multiple states of melanocyte-derived cells explain the distinct attributes of melanoma, such as invasiveness and drug resistance. Although genetic mutations can lead to drug-resistant melanoma, non-genetic mechanisms have also been attributed to the emergence of drug-resistant states^29, 30^.

The co-existence of such transcriptionally distinct NHEM sub-populations in the same environment is not known, and if they exist, whether they could be interchanged remains speculative. NHEMs can dedifferentiate to neural crest-like cells under appropriate culture conditions, revealing their potentially plastic nature^31^. Currently, melanocytes can be derived from induced pluripotent stem cells (iPSC) using multiple methods^32, 33^. A recent study showed that the melanocytes derived from temporally distinct iPSC-derived progenitors differ in their pigmentation, migration and proliferation potential^34^. Another study identified a stem cell population in the interfollicular epidermis, which can give rise to pigmented melanocytes^35^. In some altered states of the skin, such as hypertrophic scars following burn injury, amelanotic melanocytes are observed that can potentially become pigmented upon treatment with alpha melanocyte stimulating hormone (α-MSH)^36^. These studies indicate the presence of multiple states of melanocytes in the epidermis.

In this study, we employ single-cell-based methodologies to identify the transcriptional and phenotypic states of melanocytes. We demonstrate that human epidermal skin harbours multiple melanocyte states, and some of these states could be captured under culture conditions. In the temporally resolved progressive pigmentation model, we observe stochasticity in attaining the high and low-pigmented (HP and LP) state. Using scRNA seq, we demonstrate a sequential transition of the population dynamics over time and the pre-existence of the transcriptional program for the HP cell state. Moreover, we observe the coexistence of proliferation and pigmentation states across both the progressive pigmentation model and the cultured NHEMs. Differential activation of the α-MSH pathway using a pharmacological activator alters the cell state dynamics, shifting the melanocytes to either proliferative or pigmented states. A GRN derived from active enhancers in the two states independently simulates the coexistence of multiple melanocyte states. Using two independent cell-based model systems, we establish a conserved network that sustains two broad melanocyte states differing in proliferation and pigmentation.

## Results

### Human skin melanocyte pool contains distinct functional melanocyte states

To identify distinct functional states in the epidermal melanocyte pool, we reanalysed the NHEM scRNA data from GSE151091^37^. After the initial dimensionality reduction and clustering of the melanocyte subset using Seurat^38^, we observed that age group, donor ID, or plate ID were a possible source of heterogeneity that was reflected in the UMAP space (Fig S1A). While Belote et al. ^37^ identified the age and site-specific differences within the human melanocyte pool, we were interested in looking beyond and identifying functionally distinct melanocyte states, if any, irrespective of their location in the body or the age group from which they were isolated. Hence, we regressed out these effects using Harmony^39^ and identified five distinct clusters (Fig S1B), possibly representing distinct functional states of melanocytes. Cluster markers and their functional annotation revealed two mature clusters, both of which showed high expression of *MITF*, *TYR*, and *DCT*, one proliferative cluster which showed enrichment of *MKI67* and *TOP2A*, and one stem cell-like cluster enriched for genes such as *TWIST1*, *TWIST2*, and *AXL*. Interestingly, one cluster showed enrichment of interferon (IFN) signaling genes, such as *IFIT1*, *IFIT2* and *STAT1* (Fig 1A, S1C, S1D).

**Fig 1:**
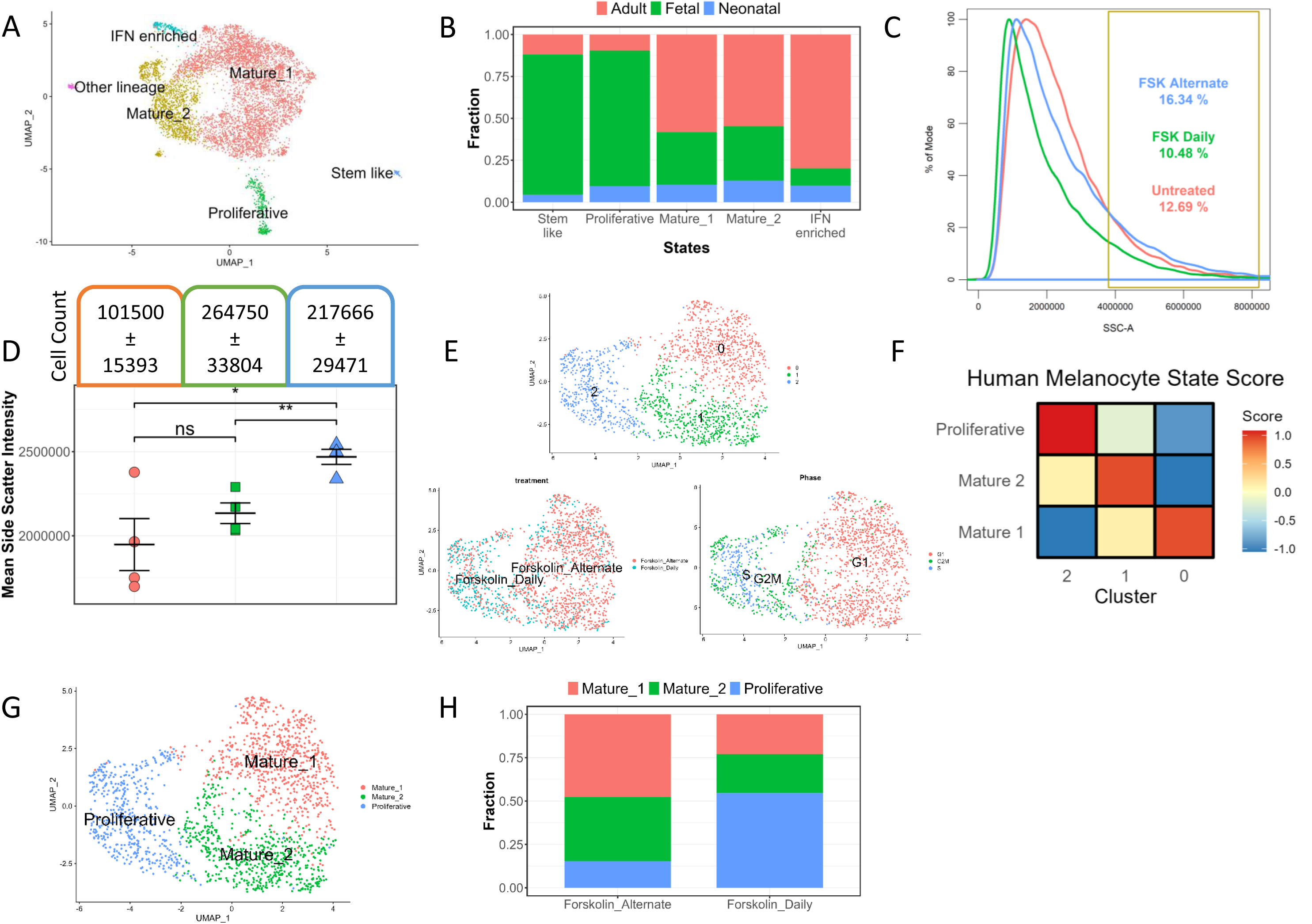
Human skin melanocyte pool contains distinct functional states. A. UMAP visualization of the human skin melanocyte colored by the identified states B. Stacked bar plot showing the distribution of different age groups in each of the identified cell states. C. Density plot showing the distribution of pigmentation in cultured NHEM with daily or alternate day treatment of forskolin. D. Mean side scatter intensity and cell count of cultured NHEM treated with daily or alternate day treatment of forskolin. (unpaired students t test, n=4, *: p value ≤ 0.05, **: p value ≤ 0.01, ns: p value > 0.05) E. UMAP visualization of cultured NHEM colored by the clusters F. Heatmap showing the scores (z-score of mean) of each of the cultured NHEM clusters for the human melanocyte mature 1, mature 2 and proliferative state. G. UMAP visualization of cultured NHEM colored by the annotated states based on cluster markers and human skin melanocyte state score H. Stacked bar plot showing the proportion of different melanocyte states upon daily and alternate forskolin treatment.

While studies have reported IFN gene signatures in melanocytes^40, 41^, especially in disease conditions such as vitiligo^42^, this is the first time a stable state enriched for interferon signatures has been identified in NHEMs. While the majority of the mature and the IFN-enriched cells came from the adult sample, the stem cell-like and the proliferative cells were enriched in the foetal skin. Although the proportions of these sub-populations were different, these states were present across all age groups (Fig 1B). This analysis corroborates Belote et al.’s ^37^ findings and suggests the existence of multiple melanocyte cell states in the human epidermal skin. The same could be observed in another publicly available human melanocyte dataset (Fig S2A-C). Therefore, it is tempting to speculate that melanocytes, much like immune cells and neurons, can exhibit distinct functional states in the same environment^43, 44^.

These transcriptionally distinct clusters are deduced from cross-sectional single-cell data sets from the human epidermis. This raises the following questions: do these sub-populations or clusters represent distinct functional states? Does each cell in these clusters transition between these states following a set trajectory? Or are these states stable and self-sustaining in nature?

To address these questions, we revisited the NHEMs in culture. We used an already established mode of altering melanocyte phenotype using forskolin (fsk) as an external agent. Fsk elicits a UV-like response in melanocytes by activating cyclic adenosine monophosphate (cAMP) and the downstream signalling cascade. Daily treatment results in a more proliferative phenotype, while treatment every alternate day gives rise to a hyperpigmentation response at the population level^45^. To dissect these phenotypes at a more granular level, we resorted to a flow cytometry-based approach to estimate pigmentation at a single-cell resolution^46^, while cell count served as a surrogate for proliferation. In the untreated melanocytes, we observed a gradient of pigmentation with a non-normal distribution containing a shoulder peak representing highly pigmented cells (Fig 1C). Daily treatment with fsk reduced while alternate day treatment increased this population (Fig 1D), suggesting that this model offers a platform to dissect these potential functional states.

We performed single-cell RNA sequencing on the daily and alternate-day fsk-treated melanocytes. Upon dimensionality reduction and clustering, we observed three clusters, with two composed of cells mostly from the alternate-day treated sample while the third contained cells mostly from the daily-treated sample (Fig 1E). Scoring these clusters for the human skin melanocyte states revealed that cluster 0 scored highest for the mature 1 state, cluster 1 for the mature 2 state, and cluster 2 for the proliferative state (Fig 1F). This, along with the cluster markers and GO term enrichment, allowed us to infer the identity of the clusters (Fig 1G). Based on the cluster composition, the daily and alternate-day treated samples contained all three identified states in different proportions. Although the stem cell-like and IFN-enriched states were not observed, probably due to non-conducive culture conditions, three of the five states namely proliferative, mature 1 and mature 2 were faithfully recapitulated in cultured NHEMs. Our observations thus confirmed that, indeed, these melanocyte states are stable and co-exist. As inferred from the flow cytometry data, alternate-day treatment enriches the mature states, whereas daily fsk treatment enriches the proliferative state (Fig 1H). This observation suggests that the distinct states of melanocytes identified in this study are also interconvertible, depending upon the external cue. Whether all states are equally responsive to the cue and how these transitions are orchestrated remains to be understood.

### Progressive pigmentation model exhibits stochastic melanocyte state transitions

To identify the core networks maintaining the melanocyte cell states and regulating the transitions between states, we leveraged the already established B16 pigmentation model that mimics the tanning and de-tanning response^47^. This model offers a temporal resolution of pigmentation induction, allowing us to capture any alterations in cell states and complement the NHEM-based approach. Though a progressive pigmentation of the cell population is observed at the bulk level in this system, heterogeneity in melanin content is observed at the microscopic level. Flow cytometry offers a way to identify pigmented cells within the population^46, 48, 49^. We used this information to develop a method to estimate melanin content in single cells using imaging flow cytometry (Fig S3A, S3B). Though at Day 7 the cell pellet is completely black with eumelanin, imaging flow cytometry revealed that the cells show a clear bimodal pigmentation distribution (Fig S3C). This confirms the presence of a heterogeneous cell population in this progressive pigmentation model, endorsing the observations made in cultured NHEMs.

NHEM-based state transitions are unlikely to be driven by genetic mutations. The progressive pigmentation model, being a cancer cell-based system, could have acquired mutations over the course of time, resulting in the observed heterogeneity. The other possibility would be that at some point during progressive pigmentation, a few cells “switch on” the expression of pigmentation genes and then are destined to become pigmented, which would suggest stochastic (non-genetic) state transitions. To rule out the genetic component, we used a single clonal cell-derived population for further experiments. Two independent clonal populations gave rise to the similar bimodal distribution observed earlier, suggesting this phenomenon to be non-genetic in nature. Since this progressive pigmentation model is a cell-density dependent phenomenon, we employed another approach to test the genetic or non-genetic nature of this heterogeneity. Herein, we subjected the cells to one round of pigmentation model and different treatments to obtain varying proportions of low and high-pigmented cells. The subsequent round of progressive pigmentation was initiated using these differentially pigmented cell populations, and the pigmentation state of the cells was assessed. We observed that irrespective of the starting proportion of pigmented cells, the final distribution of pigmentation was the same for all samples suggesting stochasticity in the system (Fig 2A). Since, in this model, cells grow as separate colonies, we assessed the colony pigmentation using high-throughput imaging. We observed that the colonies also showed the same pigmentation distribution irrespective of the starting proportion of pigmented cells (Fig 2B), thereby confirming that the transitions in this system are stochastic in nature.

**Fig 2:**
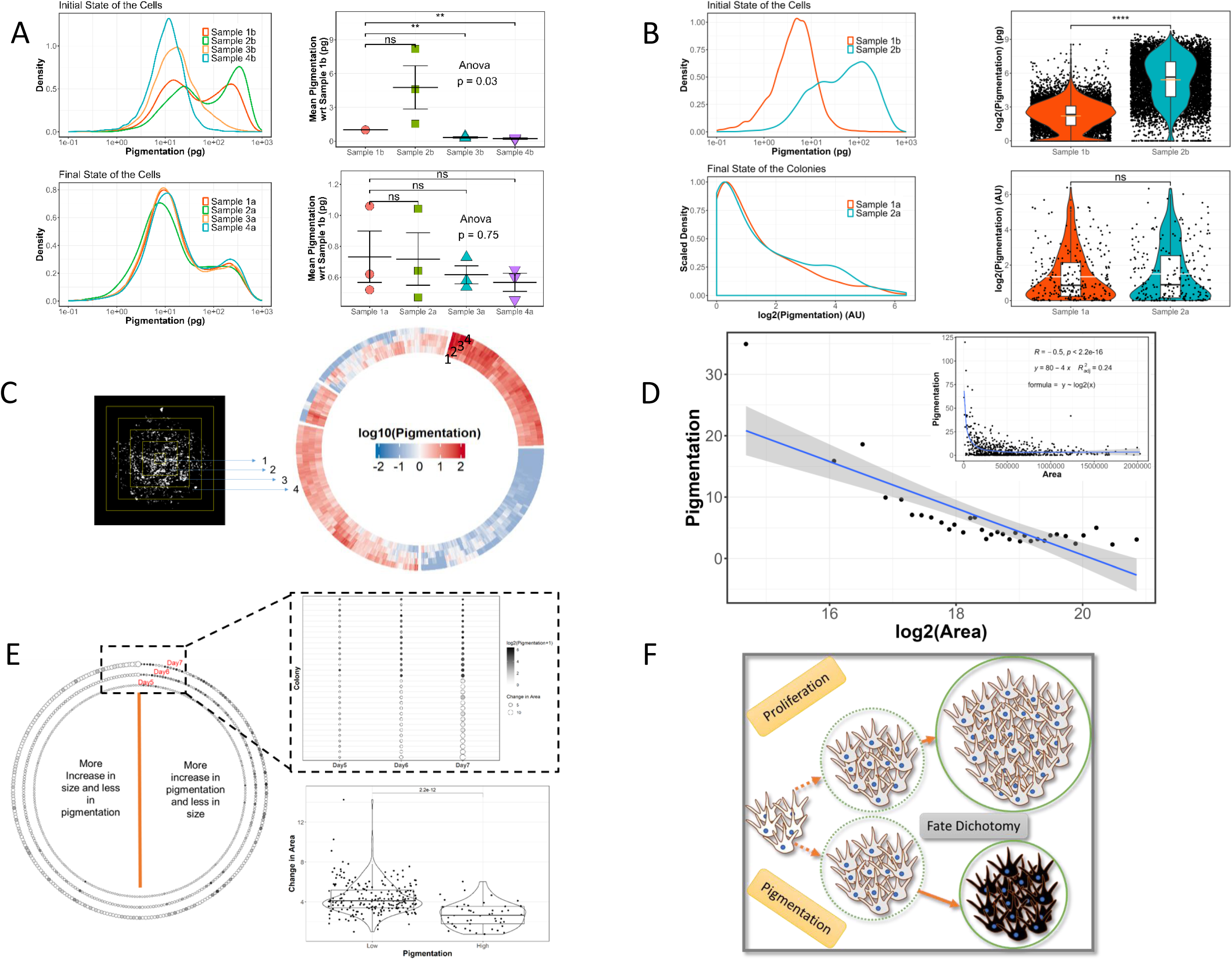
Progressive pigmentation model exhibit stochastic melanocyte state transitions. A. Density plot showing the distribution of pigmentation (log scale) in single cells across different B16 samples before (top left) and after (bottom left) seeding and comparison of mean pigmentation normalized to sample 1b before (top right) and after (bottom right) seeding (Wilcoxon-Mann-Whitney test, n=3, **: p value ≤ 0.01, ns: p value > 0.05) (Sample 1b: Untreated, Sample 2b: 50µM IBMX, Sample 3b: 100µM PTU, Sample 4b: 200µM PTU). B. Density plot showing the distribution of pigmentation (log scale) in single cells and colonies across different B16 samples before (top left) and after (bottom left) seeding respectively and comparison of mean pigmentation in single cells and colonies before (top right) and after (bottom right) seeding (Wilcoxon-Mann-Whitney test, n=3, ****: p value ≤ 0.0001, ns: p value > 0.05). C. Representation of division of colonies into concentric rectangles and the slices in between the rectangles (left) and circular heatmap (right) showing the pigmentation in each of the 4 slices (inner to outer circle) for each colony. D. Binned scatter plot showing the inverse relation between colony area and pigmentation. Inset: Raw data scatter plot with correlation coefficient derived from Pearson’s correlation analysis and the equation derived from linear regression analysis between pigmentation and log2(Area). E. Time-course tracking and analysis of colony pigmentation (colour of the circle) and area (size of the circle) from day5 to day7 of the progressive pigmentation model (left), dot plot showing the top colonies with least change in pigmentation or highest change in area (top right) and statistical analysis of change in colony area from day 5 to 7 between low and high pigmented colonies (Wilcoxon-Mann-Whitney test) F. Schematic representing the dichotomy in fate decision resulting in two broad states in the progressive pigmentation model

Interestingly, assessment of the pigmentation status within the colonies revealed that the intra-colony variability was lower than the inter-colony variability (Fig 2C). The pigmentation distribution of colonies suggested that the cells maintained two broad states with either low or very high pigmentation (Fig S2D). Since each colony is derived from a single cell, this suggests that the cells are indeed biased towards maintaining either of the two broad phenotypic states. Microscopically, we observed that the colonies in the low-pigmented state were bigger compared to the high-pigmented colonies. Pearson’s correlation analysis between the colony pigmentation and area was –0.5 suggesting a moderate negative correlation between pigmentation and colony area (Fig 2D). This is indicative of pigmenting cells to have a lower proliferation potential.

As pigmentation is a time-dependent slow process, it takes around 4-7 days before the actual phenotype presents itself and is detectable in the cell. With this, it becomes important to estimate the stage at which the pigmentation process starts, which presumably would be earlier than the phenotype is observable. A time-resolved model such as the pigmentation model confers an advantage to simultaneously infer the kinetics of both pigmentation and proliferation. We performed high throughput imaging of the colonies on days 5, 6 and 7 of the pigmentation model. We observed that pigmented colonies on day 5 maintained their high-pigmentation state till day 7 and concomitantly had a lesser increase in colony area. Whereas less-pigmented colonies on day 5 proliferated more, maintaining their low pigmentation status till day 7. Thereby, these cells commit towards either pigmentation or proliferation fate as early as day 5 in this system, suggesting an early fate biasing decision that is retained in the daughter cells (Fig 2E).

These analyses revealed that B16 cells, when put through the progressive pigmentation model, exhibit stochastic state transitions leading to two broad phenotypic states: low pigmentation (LP) and high pigmentation (HP), likely via their own separate intermediate state (Fig 2F).

### Stochastic gene expression leads to transcriptionally distinct melanocyte cell states

Visually the phenotypic heterogeneity in the pigmentation model was evident. From the time course tracking of individual colonies, it could be inferred that once the cells commit to a particular fate of either pigmentation or proliferation, their progeny retains the same state. Having ruled out the genetic factor, it is likely that epigenetic inheritance could explain the daughter cells retaining the parental state^50, 51^. To address both the transcriptomic footprint and the epigenetic landscape, we performed single-cell multi-omics sequencing across days 0, 3 and 5 of the progressive pigmentation model (Fig S4A).

Quality control was performed based on both RNA and ATAC parameters, and the RNA data was analysed first to identify the transcriptionally distinct clusters. We observed four different clusters possibly representing four different states. Colouring the UMAP based on the day revealed that three of the four clusters were predominantly composed of cells pertaining to different days of the pigmentation model. A technical replicate of day 5 was included in the analysis and both the technical replicates overlayed completely, confirming the absence of batch effects driving the clustering. We then used the weighted nearest neighbour approach in Seurat and Signac^52^ to make a joint neighbourhood graph representing both the RNA and ATAC data in the UMAP space. Although the ATAC data alone was able to explain the differences in the day 0, 3 and 5 cells, it was not enough to segregate cluster 3 from the rest. Therefore, we used the RNA data to annotate the clusters. Based on the differentially expressed genes in each cluster and GO term enrichment (Supplementary Table 1) we annotated the clusters as B16 native state enriched in both stem cell-like (*Sox10*) and proliferative signatures (*Cenpa*, *Cdk1*), differentiating state enriched with markers such as *Mitf*, *Oca2* and *Mlph*, proliferative state with high expression of *Mki67* and *Top2a* and a mature state having an enrichment of classical melanin synthesis genes such as *Tyr* and *Dct* among others (Fig 3A, 3B). We observed that the native, differentiating, and proliferative states were mainly present at day 0, 3 and 5, respectively, while the mature state emerged as early as day 3 (Fig 3C). This was surprising because visually pigmented cells could not be detected on day 3. The states observed in the progressive pigmentation model bear a resemblance to the states observed in human skin melanocytes and cultured NHEMs. This model further allowed us to temporally resolve these cell states.

**Fig 3:**
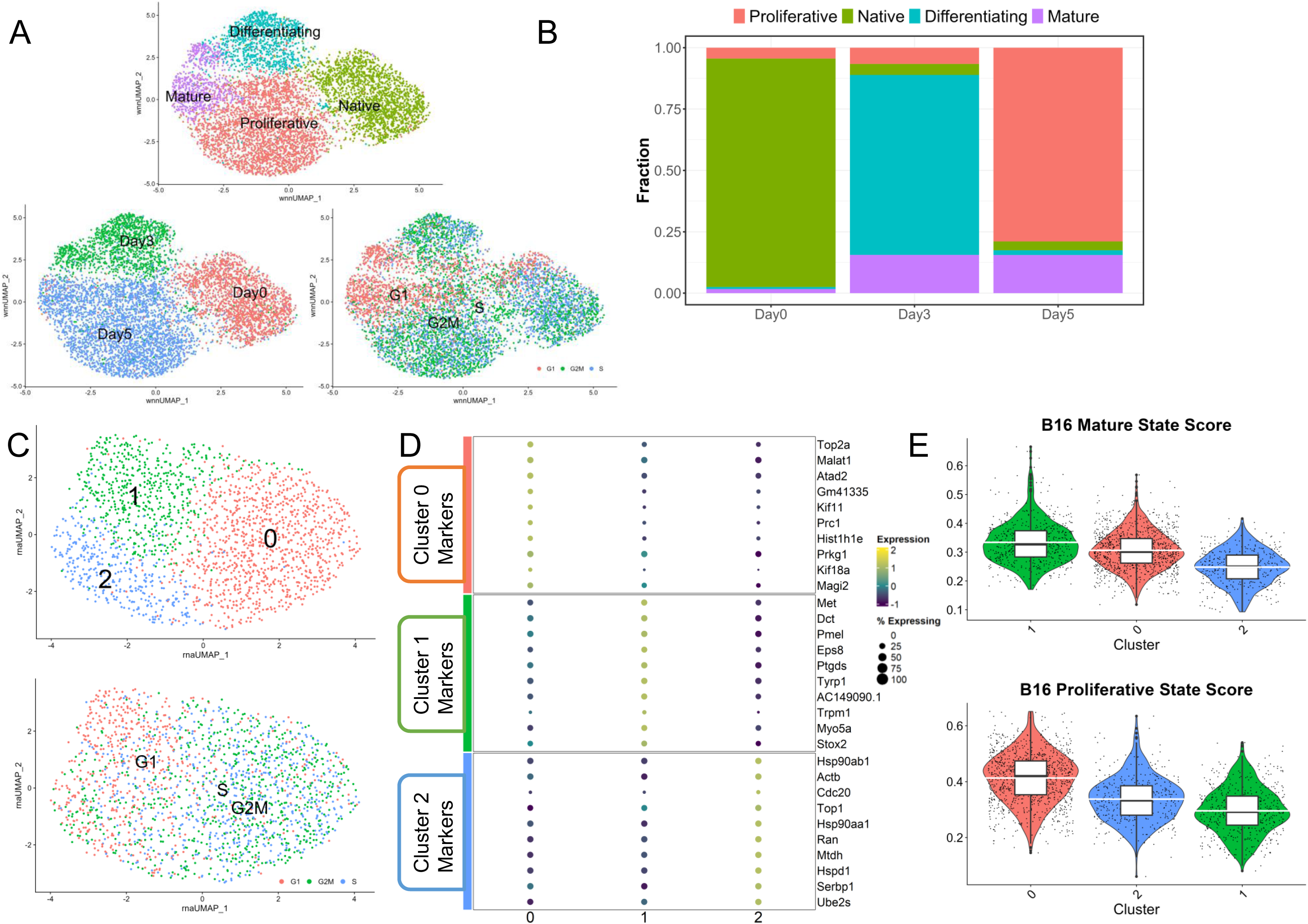
Stochastic gene expression leads to transcriptionally distinct melanocyte cell states. A. UMAP plot showing B16 cells coloured either by the clusters (top), the day of the pigmentation model (bottom left) or by their cell cycle status (bottom right) B. Stacked bar plot showing the distribution of the identified states at different days of the pigmentation model C. UMAP plot showing sub-clusters of day0 B16 cells coloured either by the clusters (top) or by their cell cycle status (bottom) D. Dot plot showing the top 15 marker genes enriched in each sub-cluster of day0 B16 cells with the size showing the percent of cell expressing the gene and colour showing the scaled mean expression value in each cluster (Wilcoxon-Mann-Whitney test with average log fold change > 0.2 and adjusted p value ≤ 0.05). E. Violin plot showing the score of each sub-cluster of day0 B16 cells for the B16 mature (top) or proliferative (bottom) state

Enrichment of different state markers in the day 0 cluster indicated the pre-existence of heterogeneity in the starting population. To investigate this further, we sub-clustered the day 0 sample individually. We used the markers of all the other states in the pigmentation model as variable features for dimensionality reduction and clustering. The cells segregated into 3 clusters with cluster 0 having relatively high expression of proliferative state markers and cluster 1 having enrichment of mature state markers (Fig 3E, 3F). Cell state scoring confirmed the relative enrichment of proliferative and mature state markers in clusters 0 and 1, respectively (Fig 3G). This confirmed our hypothesis that day 0 cells, even though a genetically and phenotypically identical clonal population, natively harbour a heterogeneous cell population, possibly due to stochastic gene expression. This heterogeneity is likely to prime the cells to switch into the proliferative or the pigmented state that are observed later in the model.

From this analysis, we confirmed that during progressive pigmentation, cells start to differentiate at day 3 and give rise to 2 broad states: proliferative and mature at the end of day 5. These sub-populations could possibly correspond to the low and high-pigmented populations we observed on day 7 using imaging-based techniques.

### Melanocyte low and high pigment states correspond to proliferative and mature transcriptional states

To map the transcriptional footprint of the low and high-pigmentation states at day 7, we sorted these two populations using a flow cytometer (Fig S4B) and subjected them to single-cell RNA sequencing. Dimensionality reduction and unsupervised clustering revealed that low and high-pigmented cells segregated into two different clusters. Interestingly, upon increasing the resolution further, two distinct sub-clusters emerged within the low and high-pigment clusters (Fig 4A). Scoring the cells for the cell cycle genes showed that both the low and high-pigmented clusters contained a G1 (non-cycling) and an S/G2/M (cycling) subpopulation (Fig 4A). Cluster markers and their GO term enrichment allowed us to infer the possible functional state of each cluster. Cluster 3 was enriched for mature state markers such as *Mitf* and *Tyr*, while cluster 2 was enriched for proliferative markers such as *Mki67* and *Top2a*. More interesting was cluster 1, containing stem cell-like markers *Klf4* and *Hes1*, but was relatively less proliferative than the other clusters suggesting a quiescent state. Cluster 0 had enrichment of both mature and proliferative markers, possibly representing a transitory state before the melanocytes acquire a hyperpigmented mature state (Fig 4B). To confirm the identity of each cluster we scored each of these for the previously identified human melanocyte state markers. Scoring confirmed our inference based on the cluster markers, with cluster 1 having the highest score for the human melanocyte stem cell-like state markers and cluster 0 having high score for both mature and proliferative states. Cluster 2 scored highest for the proliferative state, while cluster 3 for the mature state (Fig 4C). We scored these clusters for melanocyte states derived from mouse hair follicle single-cell RNA sequencing data^18^ and obtained similar results (Fig S4C).

**Fig 4:**
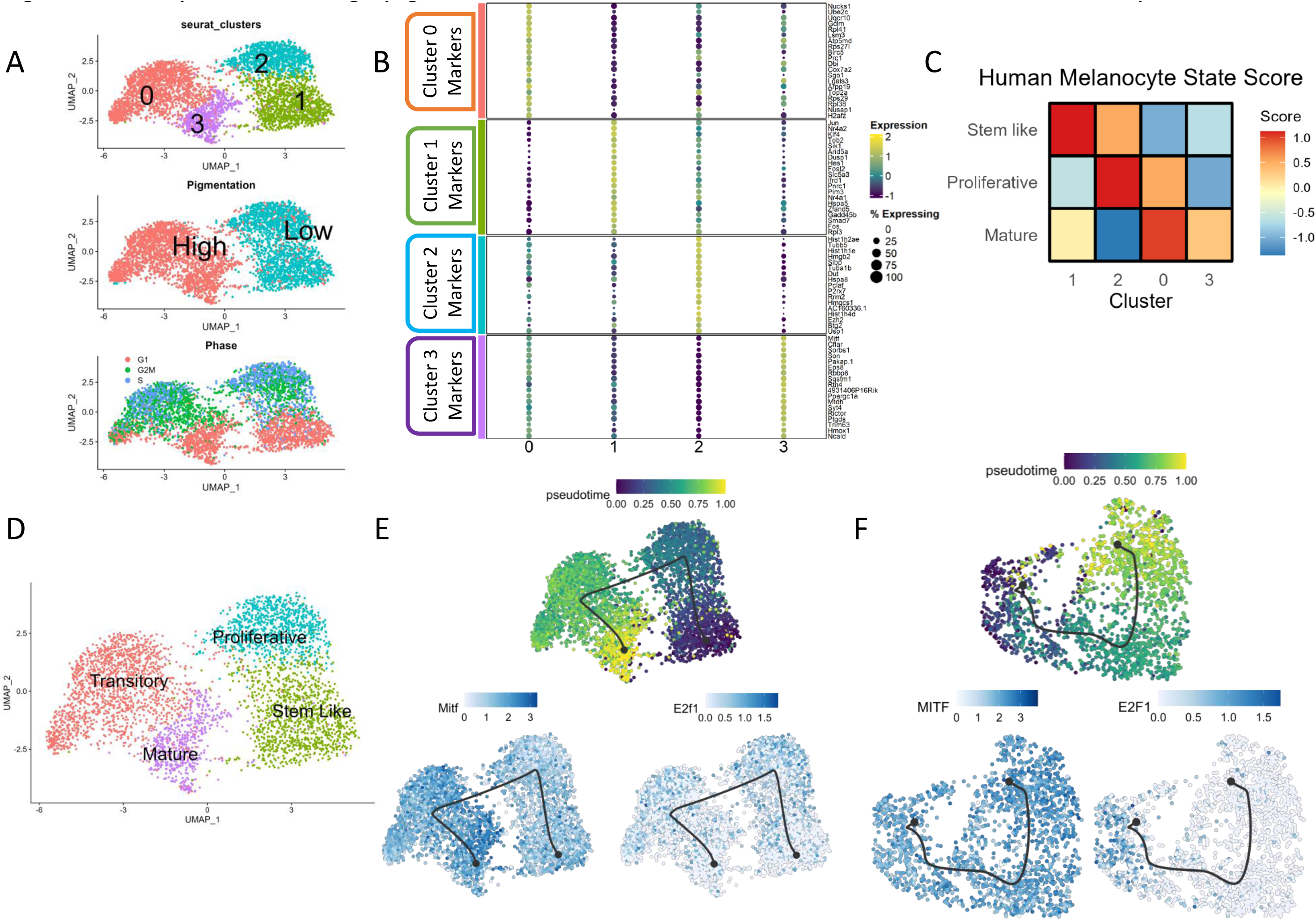
Melanocyte low and high pigment states correspond to proliferative and mature transcriptional states. A. UMAP plot showing B16 Day7 cells coloured by the clusters (top), pigmentation status (middle) or cell cycle status (bottom) B. Dot plot showing the top 20 marker genes enriched in each cluster with the size showing the percent of cell expressing the gene and colour showing the scaled mean expression value in each cluster (Wilcoxon-Mann-Whitney test with average log fold change > 0.25 and adjusted p value ≤ 0.05) C. Heatmap showing the scores (z-score of mean) of each of the B16 clusters for the human melanocyte mature, proliferative and stem cell-like state. D. UMAP plot showing B16 day 7 cells coloured by the cell states identified using marker genes, GO term enrichment analysis and scores for human melanocyte cell states. E. UMAP plot showing B16 day 7 cells coloured by pseudotime (top left), Mitf expression (bottom left) or E2f1 expression (bottom right) F. UMAP plot showing cultured NHEM coloured by pseudotime (top left), MITF expression (bottom left) or E2F1 expression (bottom right)

These results allowed us to annotate the B16 cell states (Fig 4E) and revealed the functional similarities between the B16 mouse melanoma cells and normal human skin melanocytes, endorsing the use of the B16 pigmentation model to uncover the underlying gene regulatory networks (GRNs) governing melanocyte states. It was interesting to observe that all these states were maintained even at this stage in the pigmentation model when presumably the cells would have attained their terminal state of differentiation. Whether these states could switch from one to another was still unclear. To test this *in silico*, we used the Dynverse^53^ package in R to infer a trajectory that would indicate the transition potential of these states. Upon applying the Scorpius algorithm, we observed a linear trajectory starting from the stem cell-like state and ending at the mature state with the proliferative and the transitory states in between (Fig 4F). This highlights the switching potential of the stem cell-like state to the mature hyper-pigmenting state if the need arises. A similar inference could be derived from cultured NHEM single-cell data wherein the cells transit from a proliferative to a mature state *via* an intermediate state in between (Fig 4E). This explains the different proportions of cell states observed with fsk treatments. Thereby, using two different cultured cell-based models, we demonstrate the co-existence of multiple melanocyte states with the possibility of state transitions when acted upon by an external cue.

### Active enhancers guide the melanocytes towards the mature state

While the cultured NHEM model revealed the co-existence and the switching potential of the melanocyte states, the B16 pigmentation model allowed us to trace the dynamics of these state transitions. The time-course tracking of colonies revealed that the daughter cells retain the parental states of low and high-pigmentation, pointing towards an epigenetic driving force. Epigenetics influence the cell state dynamics and possibly regulates the cell state heritability across multiple cell divisions^50, 51, 54, 55^. To identify the key players governing the identified melanocyte states, we resorted to an integrated approach involving chromatin accessibility (scATAC seq), an appropriate upstream histone modification (ChIP seq) and the downstream gene expression footprint across the LP and HP population (scRNA seq).

We analysed the scATAC seq data of the single-cell multi-omics experiment performed on days 0, 3 and 5 of the pigmentation model. We observed enrichment of open chromatin regions around the TSS for all the four states (Fig S4D). We found subtle differences in chromatin accessibility with only 2713 differentially accessible regions (DARs) (adjusted p-value ≤ 0.05) across the four states, with 371 regions enriched in the mature state (Supplementary Table 1). Melanocytes maintained two broad stable states, proliferative and mature corresponding to the low and high pigment phenotype. We compared these two states for chromatin accessibility differences and observed 397 and 162 regions to be differentially accessible in these respective states (adjusted p-value ≤ 0.05). Mapping the differentially accessible regions to the nearby genes revealed a few regions associated with known pigmentation genes such as Mitf and Slc24a5. Motif enrichment analysis yielded several transcription factor motifs among these DARs. Since single-cell ATAC seq data is very sparse in nature, which may make it difficult to capture subtle differences in chromatin accessibility across sub-states of the same cell type, accounting for the fewer DARs observed.

H3K27ac has been shown to be abundant on pigmentation genes in melanocytes^45, 56^. In one of our earlier studies, we have shown that, through a pH-mediated feed-forward loop, H3K27ac selectively activates pigmentation genes and accentuates melanocyte differentiation at the population level^57^. Hence, we decided to investigate whether H3K27ac differences could explain the existence of these cell states. As the epigenetic regulation is likely to precede the transcriptional differences observed on day 7, we selected day 5 for our analysis. We performed chromatin immunoprecipitation of H3K27ac followed by sequencing (ChIP seq) in LP and HP cells sorted using flow cytometry on day 5 of the pigmentation model (Fig S4E). Peak calling, using a well-established program, macs3^58^, gave us 13008 peaks in the LP, while 25999 peaks in the HP, a marked increase compared to LP. Of these, 14368 were unique to HP, while only 723 were unique to LP. Annotating the peaks using the ChIPseeker package^59^ in R revealed that around 80% of the marked regions were within the 3kb promoter in LP, which reduced to around 50% in HP with a concomitant increase in the first intron and distal intergenic region (Fig S5A). Differential peak analysis using DiffBind^60^ gave us 12579 differentially marked regions (FDR ≤ 5%), with almost all of those (12540) having increased H3K27ac in HP (Fig 5A). After associating these differentially marked regions with the nearby genes using the rGREAT package^61^, we observed Mitf and some of its targets to have higher H3K27ac in the HP population. Motif enrichment analysis using HOMER^62^ revealed 242 motifs, corresponding to several transcription factors (TFs) including MITF and Lef1, in the differentially hyperacetylated regions (DHAcRs) of HP.

**Fig 5:**
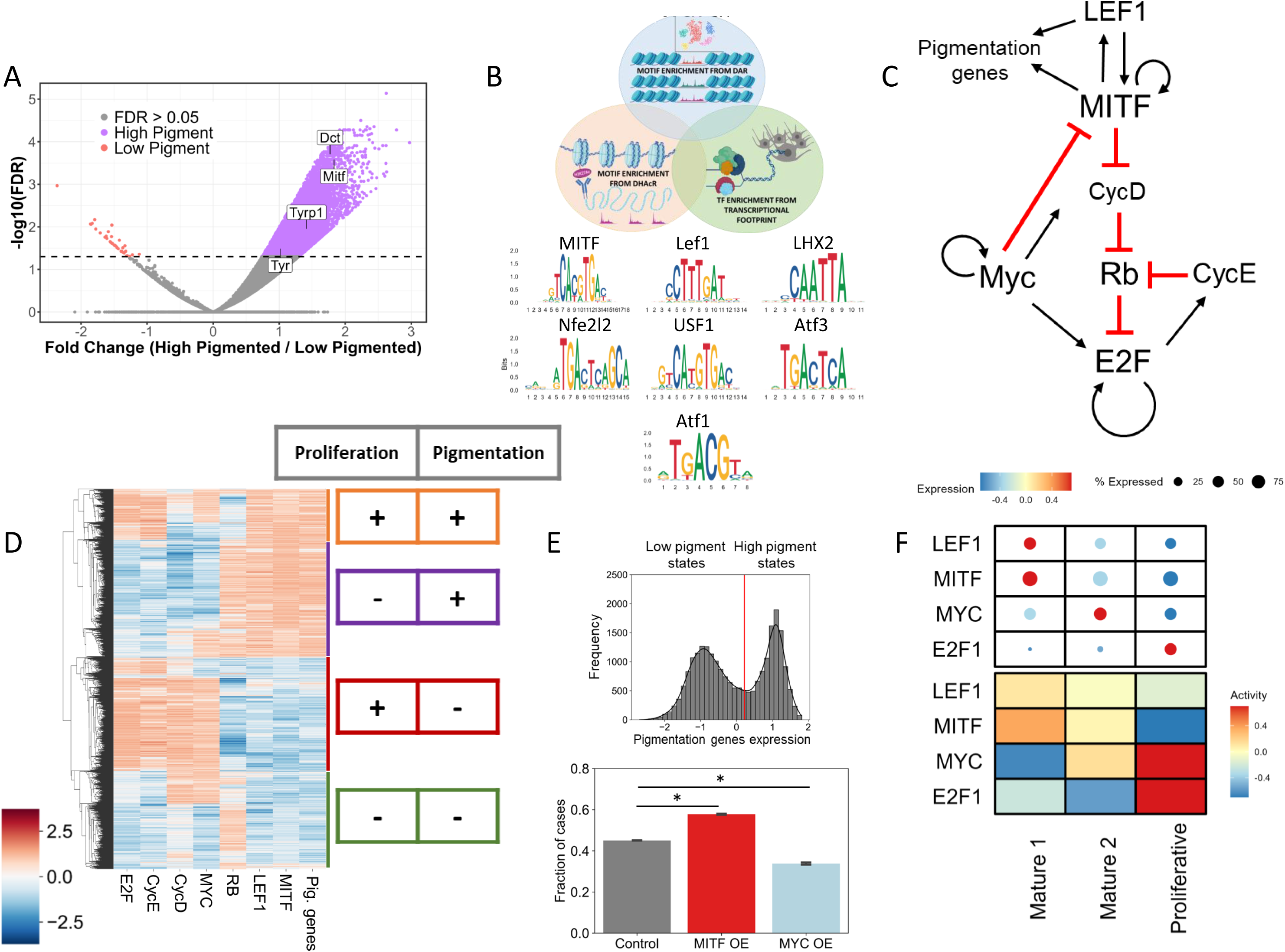
Active enhancers guide the melanocytes towards the mature state. A. Volcano plot showing regions differentially acetylated at H3K27 in day 5 LP and HP population highlighting some of the pigmentation genes. B. Motif plot showing motifs derived from the intersection of motifs enriched in DHAcR in day 5 HP, DAR in mature state and TF’s enriched in day 7 HP population. C. Gene regulatory network, constructed from the common motifs, responsible for governing melanocyte cell state transitions. D. Steady state solutions of RACIPE simulations showing the presence of various stable states allowed by the gene regulatory network. Colour bar shows the z-normalised simulated expression levels of the listed genes with red representing higher expression and blue representing lower expression. E. Histogram with corresponding kernel density estimate showing the bimodality in the expression of pigmentation genes (top) and simulation results showing effects of perturbation on the frequency of high pigmented states upon MYC over expression and MITF overexpression (students T-test, *: p value ≤ 0.01) F. Heatmap showing expression (top) and activity (bottom) of TF’s governing melanocyte cell states in cultured NHEM single-cell RNA seq data.

In the third approach, we set out to identify TF activity that could explain the differential gene expression across LP and HP states on day 7. We used Dorothea and Viper libraries in R^63–65^, to estimate the TF activity from the gene expression data. After constructing the TF activity by cell count matrix, we performed unsupervised clustering and observed that LP and HP segregated into two different clusters. This allowed us to identify the TFs that govern the differential transcriptome in the LP and HP population (Fig S5B). We then identified the common overlapping set of TFs whose footprint is represented in all the three independent analyses described above. This resulted in a set of 7 TFs enriched in HP (Fig 5B). A similar analysis for LP population resulted in a larger set of 86 common TFs, as there were no DHAcRs in LP (Supplementary Table 1).

### Melanocyte cell state GRN reliably predicts the state transitions

To construct a GRN that explains the coexistence and transitions of melanocyte cell states, we used the state-specific TFs identified above along with evidence from the literature^66–70^. Cell cycle as a bistable switch has been modelled previously in the context of general mammalian cell cycle entry decision making circuit^71^. We used this information and constructed a GRN coupling pigmentation associated genes – MITF and LEF1 with the cell cycle associated genes – RB, MYC, CycE and CycD (Fig 5C).

Next, we sought to simulate it to study the steady states allowed by this model and if it can explain the inverse association between melanocyte pigmentation and proliferation. To achieve this, we used the computational framework RACIPE (Random Circuit Perturbation), which solves a set of coupled ordinary differential equations (ODEs) to examine the various phenotypic states enabled by a GRN^72^. It does this by sampling an ensemble of kinetic parameter sets from a biologically relevant parameter range. For each distinct parameter set, it chooses initial conditions based on random sampling from within a log-uniform distribution for each node and then solves ODEs to obtain the possible steady states^73^. For some parameter sets, more than one steady state (phenotype) is achieved, suggesting possible stochastic switching among those states under the influence of noise.

Using RACIPE, we found that the constructed GRN can give rise to four main states – a high-pigmented state with low cell cycle genes, a state expressing both pigmentation and cell cycle genes, a high proliferation state with low pigmentation genes and a non-cycling stem cell-like state (Fig 5D). Simulated expression profile of pigmentation genes showed a clear bimodal distribution with distinct high and low pigmentation states (Fig 5D). Principle component analysis highlights the expression of MYC, E2F, and pigmentation genes (Fig S5D), unravelling four states similar to those observed in our single-cell data (Fig 4D).

Mitf is the master regulator of melanocyte biology, associated with multiple functions^74^. An increase in Mitf levels is suggested to control melanocyte differentiation and survival, and its expression relatively goes down with pigmentation^45, 75^. Our multi-omics experiment suggested a similar scenario with the differentiating state (observed on day 3) showing the highest *Mitf* expression that decreased subsequently on day 5 (Fig S5E). One of the consistent themes that emerged from all our analyses was that in already differentiated melanocytes, *Mitf* had significantly high expression in the mature state compared to the proliferative state across all the datasets (Fig S5F). Therefore, we simulated the overexpression of MITF or MYC and investigated the effects on the frequency of high-pigmented state. While overexpression of MITF significantly increased the proportion of high-pigmented state, overexpression of MYC decreased it, thus pushing the cells towards a high-proliferative state. Overall, the results suggest that the steady-state dynamics of the underlying GRN play a crucial role in the emergent property of inverse association between melanocyte pigmentation and proliferation programs.

### State switching in cultured NHEMs is guided by the melanocyte cell state GRN

We revisited the forskolin response in melanocytes that activates either proliferation or pigmentation programs based on the frequency of administration (Fig 1H). Our scRNA seq analysis demonstrated that the treatments rather altered the proportion of the cell states enriched in these two diametrically opposite programs within the population. The GRN derived from the B16 progressive pigmentation model explained the co-existence of distinct B16 cell states. We then tested whether this GRN could explain the co-existence of proliferation and pigmentation states in NHEMs. We derived the activity of the key TFs in this GRN using the target gene expression in the proliferative and the two pigmenting states (mature 1 and 2). We observed that LEF1 and MITF were active in the mature 1 state, whereas MYC and E2F1 were active in the proliferative state (Fig 5E), suggesting that the same GRN is responsible for the co-existence and transitions of the human melanocyte states.

## Discussion

Here, using *in vitro* cell-based models and single-cell methodologies, we showed that specified melanocytes exhibit distinct cell states that are governed by a gene regulatory network containing regulators of pigmentation (MITF, LEF1) and proliferation (MYC, E2F1). We coupled the known proliferation circuit with the pigmentation regulators and showed that variability in the expression/activity of these factors, under the influence of noise (stochasticity), resulted in the co-existence of multiple functionally distinct melanocyte states. Physiologically, during the hair cycle, melanocyte stem cells are known to differentiate to give rise to a mature pigmented state through a proliferative state. This is shown to be governed by the interplay of external signalling factors that are spatiotemporally controlled^18^. A recent study showed that the transit amplifying (proliferative) state can either differentiate into a mature pigmented state or dedifferentiate into an McSc state in a WNT-dependent manner ^19^. Whether it is the cue that primarily induces these states, or these are pre-existing within the seemingly homogeneous melanocyte population remained unclear.

Stochasticity in gene expression has been shown to govern melanoma cell states, giving rise to a drug-resistant state even before the addition of any drug. These states are largely predetermined and are further strengthened by epigenetic programs resulting in clonal expansion^9, 30, 51^. In this current work, we observed pre-existing transcriptional variability in day 0 cells with a mature-like and a proliferative sub-population (Fig 3C-E). It is likely that these cells are the ones that give rise to the mature state in the progressive pigmentation model, and low-density culturing appears to be the trigger for these state transitions. Using high-throughput imaging on days 5, 6 and 7 in the model, we observed that the two broad states, LP and HP, are maintained throughout as independent colonies suggesting inheritance of the parental state across subsequent cell cycles. Our earlier work demonstrated the role of H3K27ac in melanocyte differentiation through a pH-mediated feed-forward loop involving Mitf and its targets. We observed hyperacetylation of pigmentation genes on day 5^57^. In this work, we map the hyperacetylation of H3K27 predominantly to the HP state compared to the LP state. This suggests that the mature and the proliferative state, much like the drug-resistant state in melanoma, are predetermined, with the mature state getting reinforced via hyperacetylation at H3K27 whenever a trigger is presented.

While the existence of proliferative and mature states is understandable, we also identified a sub-population of melanocytes having a characteristic IFN signalling gene signature. IFN-γ secreting CD8+ T cells are known to play a role in vitiligo pathogenesis leading to melanocyte destruction, and melanocytes from vitiligo skin have been shown to harbour a footprint of IFN-stimulated genes^42^. An IFN signature enriched melanocyte state in healthy individuals may represent a vulnerable melanocyte sub-population.

In a normal physiological scenario, melanocytes are constantly exposed to UV radiations resulting in the tanning response. While the response of cells to fsk is well recognized, an elegant study by Malcov-brog et al. ^45^ demonstrated that proliferation and pigmentation are differentially activated depending upon the frequency of exposure. Our study, using *in vitro* cell models, deciphers that melanocytes exhibit these two programs in separate sub-populations that pre-exist. Differential exposure to fsk results in the enrichment of one state. On recurrent exposure, the proliferating state is expanded, while infrequent exposure leads to the enrichment of the pigmentated state. Mitf dynamics remain at the centre of melanocyte functioning. Malcov-brog et al. ^45^ demonstrated that an increase in Mitf levels induces the cell survival program in the melanocyte population. In the progressive pigmentation model, we observed the highest *Mitf* levels in the differentiating state, followed by a decrease once the cells differentiate (Fig S5D). Interestingly after differentiation, we observed that cells in a proliferative state had relatively lower *Mitf* expression compared to the mature population (Fig S5E). Drawing parallels from this model suggests that exposure of melanocytes to a cue such as UV radiation/fsk results in the elevation of Mitf, leading to the initiation of the cell survival program and increased proliferation. Subsequently, Mitf decreases, and in the absence of additional exposure, cells with relatively higher levels of Mitf get biased towards the mature state while others remain in the proliferative state. The proliferation regulators such as Myc then further suppress Mitf to maintain the other cells in the proliferative state. However, further investigation is needed to delineate these cell state dynamics in detail. In all, stochasticity coupled with a programmed regulatory network explains the co-existence of melanocyte cell states, and ultimately, the population-level diversity. The co-existence of these states is likely to permit the system to adapt quickly to external cues by altering the cell state dynamics.

## Supporting information

This file contains supplementary Figures S1 to S5

## Acknowledgements

We thank Partha Chattopadhyay, Shweta Sahni and Pooja Sharma for help with single cell sequencing.

This project was funded by Council of Scientific and Industrial Research (CSIR) through the project Regen-X (MLP2008).

AA acknowledges CSIR for the support of his fellowship.

## Author Contribution

Conceptualization, VTN, RSG and AA; Methodology, AA, VTN, SS, MKJ, RP, MF and AS; Investigation, AA, VTN and AN; Writing, review and Editing, AA and VTN; Visualization, AA, AN and SS; Funding acquisition, VTN; Resources, VTN, RP and MF; Supervision, VTN, AS and MKJ

## STAR Methods

### B16 pigmentation model and NHEM culturing

All the cells were maintained at 37°C and 5% CO_2_.

B16 cells were cultured in Dulbecco’s modified Eagle’s Media (DMEM-high glucose, Sigma-Aldrich, D5648) supplemented with 10% foetal bovine serum (FBS, Gibco, 10270106). For the progressive pigmentation model, B16 cells were seeded at a very low density of 100 cells/cm^2^ and cultured for appropriate number of days as per the experimental requirement, for a maximum of 7 days. As per the requirement, the cells were either treated with 50 µM 3-Isobutyl-1-methylxanthine (IBMX, Sigma-Aldrich, I7018-100MG) to enhance pigmentation or with 100/200 µM N-Phenylthiourea (PTU, Sigma-Aldrich, P7629) to prevent pigmentation, one day after seeding the cells.

NHEMs were purchased from Lonza (CC-2504). Cells were revived in Lonza MGM4 media (CC-3250) supplemented with supplied factors (CC-4435). Post revival, cells were transferred to and maintained in Medium 254 (Gibco M254500) supplemented with human melanocyte growth supplement (S0025). 50,000 cells were seeded in 3 wells of a 6 well plate. One well was treated daily while the other was treated every alternate day with 20 µM forskolin (Sigma-Aldrich, F3917) for 7 days. Equal volume of DMSO was added in the remaining well to serve as a control.

### Melanin estimation using NaOH method

Melanin estimation was performed as described earlier^76^. Briefly, equal number of cells (∼25 Lacs) from each sample were centrifuged at high speed, dissolved in 250ul of 1N NaOH, vortexed and kept at 80°C for 2 hrs. The solution was spun down at high speed and 100ul supernatant, in duplicates, was transferred to a 96 well plate. The absorbance was measured at 400nm. Synthetic melanin (Sigma-Aldrich, M8631) was used to generate the standard curve and estimate the melanin content in each sample.

### Melanin estimation using Imaging flow cytometer

Cells were trypsinized, washed with 1x Dulbecco’s phosphate buffer saline (DBPS) two times and resuspended in 1x DPBS. Each sample was run sequentially on Amnis Imagestream Mk II after initializing the system. Initial gating was done using the area and aspect ratio parameters to select only single round cells. 10,000 cells were captured for each sample after the gating. The analysis was done on the Ideas software.

### Pigmentation estimation using ImageJ

All the microscopy images were analysed in Fiji. A threshold was applied to the images such that only the dark spots corresponding to melanin were visible in pigmented colonies and nothing was visible in depigmented colonies. Colonies were marked using the ROI tool and mean grey value estimation for each colony was taken as the pigmentation content in the colonies. Any colonies appearing to be merged (morphologically) from multiple individual colonies were left out of the analysis.

### Time-course tracking and analysis of B16 colonies

High content live cell imaging was done at day5, 6 and 7 of the pigmentation model. First the colonies at day 5 were marked using the ROI tool and then the same colonies were traced at day 6 and then at day 7 by using the edges of the well as reference. Colonies that got merged and were untraceable in the subsequent days were filtered out. Colony pigmentation and area was estimated using Fiji and the data was plotted in R Studio using ggplot2.

### Human melanocyte single-cell data analysis

The human melanocyte single-cell RNA sequencing data was analysed using Seurat v4 in R Studio. Raw counts were downloaded from GSE151091. The data was filtered for low quality cells and gene count was normalized. Clustering and dimensionality reduction was performed using first 29 PC’s. Melanocytes were extracted based on the gene expression profile and were further sub-clustered. The clusters were batch corrected using harmony.

### Single cell RNA sequencing and analysis of B16 cells

B16 cells were taken at Day7 of the pigmentation model and sorted into low and high pigmented population using side scatter intensity. The single-cell RNA libraries were prepared for both samples using the 10x genomics Chromium Next GEM Single Cell 3’ Reagent Kit v3.1. The library QC and quantification was done using Agilent bioanalyzer HS DNA kit. The libraries were pooled and sequenced on the NextSeq 2000 platform. Raw bcl files were converted to final count matrix using Cell Ranger v6.1.2 software following the tutorial provided on the 10x genomics website. All the downstream analysis was performed in R Studio using Seurat v4 package. Low quality cells and genes expressing in less than 3 cells were filtered out. The data was log normalized and 3000 highly variable genes and 30 principal components were used for dimensional reduction and clustering.

### Single cell multi-omics sequencing and analysis of B16 cells

B16 pigmentation model was setup on different days to obtain Day 3 and Day 5 sample on the same day and the cells used to seed Day 3 and 5 sample were maintained at high density and taken as Day 0 sample. The multiome libraries were prepared using the Chromium Next GEM Single Cell Multiome ATAC + Gene Expression kit. The library QC and quantification was done using Agilent bioanalyzer HS DNA kit. The libraries were pooled and sequenced on the NextSeq 2000 platform. Raw bcl files were converted to final count matrix using Cell Ranger ARC v2.0.2 software following the tutorial provided on the 10x genomics website. All the downstream analysis was done in R Studio using Seurat v4 and Signac v1.9 packages. Both RNA and ATAC parameters were used for initial quality control. The clusters were identified using the RNA data and the ATAC data was overlayed on it using joint neighbour calling to finally arrive at the joint UMAP reduction.

### Single-cell sequencing and analysis of NHEMs

Forskolin daily and alternate treated cells were taken for library preparation using the 10x genomics Chromium Next GEM Single Cell 3’ Reagent Kit v3.1. The single-cell RNA libraries were prepared for both samples using the 10x genomics Chromium Next GEM Single Cell 3’ Reagent Kit v3.1. The library QC and quantification was done using Agilent bioanalyzer HS DNA kit. The libraries were pooled and sequenced on the NextSeq 2000 platform. Raw bcl files were converted to final count matrix using Cell Ranger v6.1.2 software following the tutorial provided on the 10x genomics website. All the downstream analysis was performed in R Studio using Seurat v4 package. Low quality cells and genes expressing in less than 3 cells were filtered out. The data was log normalized and 3000 highly variable genes and 22 principal components were used for dimensional reduction and clustering.

### H3K27ac Chromatin Immunoprecipitation and Sequencing

Chromatin immunoprecipitation was performed using the protocol provided by Upstate Biotechnology with modifications as suggested in the Fast ChIP protocol. Briefly, B16 cells were taken at Day 5 of the pigmentation model. Cells were sorted into low and high pigment population based on side scatter intensity, fixed with 10% formaldehyde (Sigma-Aldrich, F8775) at 37°C for 10 mins and resuspended in SDS lysis buffer (1% SDS (Sigma-Aldrich, L3771), 10 mM EDTA (Sigma-Aldrich, E6758), 50 mM TRIS (Sigma-Aldrich, T6066) (pH 8.1)) after washing with ice cold 1x DPBS containing protease inhibitors. Lysis was done for 30 mins on ice and then chromatin lysate was obtained by manual shearing in a bath sonicator using 6-7 cycles of 7.5 mins keeping the sonicator on for 30 sec and off for 45 sec. Chromatin was quantified using qubit HS DNA kit and equal amount of chromatin was taken for each set of replicates for chromatin immunoprecipitation using the anti-H3K27ac antibody (ab4729). Unsorted cells were used for control ChIP using anti-IgG antibody (ab172730). Libraries were prepared using NEB Ultra II DNA library prep kit (E7103S) following manufacturer’s instructions and sequencing was done, after pooling the libraries together, on the NextSeq 2000 platform.

### ChIP-seq analysis

All the analysis was done using default parameters unless specified. The raw bcl files were converted to fastq using bcl2fastq. Quality control was done using fastqc and bad quality bases were trimmed using trimmomatic (CROP:150 LEADING:3 TRAILING:3 SLIDINGWINDOW:4:15 MINLEN:50). Poly-G/poly-x overrepresented sequences and any remaining adapter traces were removed using fastp. Alignment was done to mm10 genome using bowtie2 and samtools was used to fix mate coordinates, sorting and marking duplicates to get the final bam file. MACS3 was used to call narrow peaks and bedtools was used to identify the common and unique peaks between the samples. Differentially acetylated regions were identified using DiffBind and rGreat was used to annotate the regions to the nearby genes in R Studio. HOMER was used for motif enrichment analysis in the differentially acetylated regions.

### Gene Regulatory Network Simulations

Random Circuit Perturbation (RACIPE) is a computational framework that generates an ensemble of kinetic models for a given gene regulatory network and simulates its dynamics for a range of biologically relevant parameters and initial conditions. The RACIPE input network consists of inhibitory and activating links between each node. To calculate the expression of each node in the network, RACIPE uses a set of Ordinary Differential Equations (ODEs) defined as follows: dXi/dt=gXi∏jHs(Xj,Xji0,nji,λji)−kXiXi

In the RACIPE framework, each node in the input gene regulatory network is represented by a concentration variable Xi, where i∈{1,2,3,4,5}. The expression level of each node is determined by a set of ordinary differential equations (ODEs) that consider various parameters, including the basal production rate g, basal degradation rate k, and the shifted Hill function Hs. The shifted Hill function is used to incorporate the activating or inhibitory links between nodes and to determine the production rate for each node. The parameters λ, n, and X0 correspond to each regulatory link in the network and represent the fold-change parameter, Hill’s coefficient, and threshold value for the Hill’s function, respectively. Through the RACIPE framework, an ensemble of kinetic models is generated for the given gene regulatory network, and its dynamics are simulated for a range of biologically relevant parameters and initial conditions. Initial condition for each node is randomly sampled from a log-uniform distribution of minimum to maximum levels of that node. RACIPE steady states obtained are z-normalized for all further analysis. The perturbation experiments were done by increasing the basal production rates by 20-fold and comparing the percent of high pigmented states (z-score > 0.22 obtained from the bimodal pigmentation score distribution for the control case) to the control case where the basal production rate was not over expressed.

### Statistical Analysis and graphs

All the statistical analysis was performed in R Studio using ggpubr library. All the plots were made using ggplot2 library. P value (P) > 0.05 is represented as ns, P ≤ 0.05 as *, P ≤ 0.01 as **, P ≤ 0.001 as *** and P ≤ 0.0001 as ****.

## Supplemental Information titles and legends

**Fig S1: Analysis of epidermal NHEM scRNA seq data taken from GSE151091 (related to Fig 1)**

A. UMAP visualization of the human skin melanocyte coloured by age group (top), donor ID (bottom left), plate ID (bottom right)

B. UMAP visualization of the human skin melanocyte coloured by the clusters

C. Dot plot of showing the top 10 marker genes enriched in each cluster with the size showing the percent of cell expressing the gene and colour showing the scaled mean expression value in each cluster (Wilcoxon-Mann-Whitney test with average log fold change > 0.25 and adjusted p value ≤ 0.05).

D. Dot plot showing top 5 GO terms enriched in each cluster identified using cluster markers

**Fig S2: Analysis of Analysis of epidermal NHEM scRNA seq data taken from EGAS00001002927 (related to Fig 1)**

A. UMAP visualization of the human skin melanocyte coloured by the clusters

B. Dot plot of showing the top 10 marker genes enriched in each cluster with the size showing the percent of cell expressing the gene and colour showing the scaled mean expression value in each cluster (Wilcoxon-Mann-Whitney test with average log fold change > 0.25 and adjusted p value ≤ 0.05).

C. Dot plot showing top 5 GO terms enriched in each cluster identified using cluster markers

**Fig S3: Imaging based assessment of pigmentation (related to Fig 2)**

A. Pearson’s correlation analysis between the average melanin content per cell estimated using NaOH method and the different parameters of imaging flow cytometer.

B. Linear regression analysis of mean brightfield intensity of imaging flow cytometer and average melanin content per cell estimated using NaOH method.

C. Distribution of pigmentation in B16 cells at day 7 of the progressive pigmentation model.

D. Distribution of pigmentation in B16 colonies at day 7 of the progressive pigmentation model.

**Fig S4: Quality control of different single-cell data and scoring for mouse melanocyte states (related to Fig 3 and 4)**

A. Side scatter intensity distribution (representing pigmentation) of B16 cells at day0, 3 and 5 of the progressive pigmentation model. Rectangle gate represents high pigment cells.

B. Side scatter intensity distribution (representing pigmentation) of B16 cells at day0 and 7 of the progressive pigmentation model. Rectangle gate represents high pigment cells.

C. Heatmap showing the scores (z-score of mean) of each of the B16 clusters for the mouse hair follicle melanocyte mature, proliferative and stem like state.

D. Distribution of ATAC peaks around the TSS in different melanocyte states of the progressive pigmentation model.

E. Side scatter intensity distribution (representing pigmentation) of B16 cells at day0 and 5 of the progressive pigmentation model. Rectangle gate represents high pigment cells.

**Fig S5: Analysis of H3K27ac ChIP and scRNA seq datasets**

A. Plots showing distribution of H3K27ac peaks around the TSS (top), annotation of peaks (bottom left) and distribution of peaks relative to TSS in day5 LP and HP population.

B. UMAP plot showing day7 cells of the progressive pigmentation model colored by clusters identified using TF activity (top) and pigmentation (bottom). Heatmap showing top TF active in LP and HP population at day7 (right).

C. PCA plot showing expression of Myc (top), E2f (middle) and pigmentation genes (bottom) in the artificial cells simulated using RACIPE.

D. Violin plot showing Mitf expression in differentiating, mature, native and proliferative melanocyte states in the progressive pigmentation model.

E. Violin plot showing Mitf expression in mature and proliferative states across different scRNA seq datasets: human epidermal melanocytes (top), day7 of the progressive pigmentation model (middle) and day0, 3, 5 of the progressive pigmentation model.

