## Supplementary figures and images for "Melanocytes Exhibit Distinct Cell States Governed by A Gene Regulatory Network Under Stochastic Influence"

### This file contains supplementary Figures S1 to S5

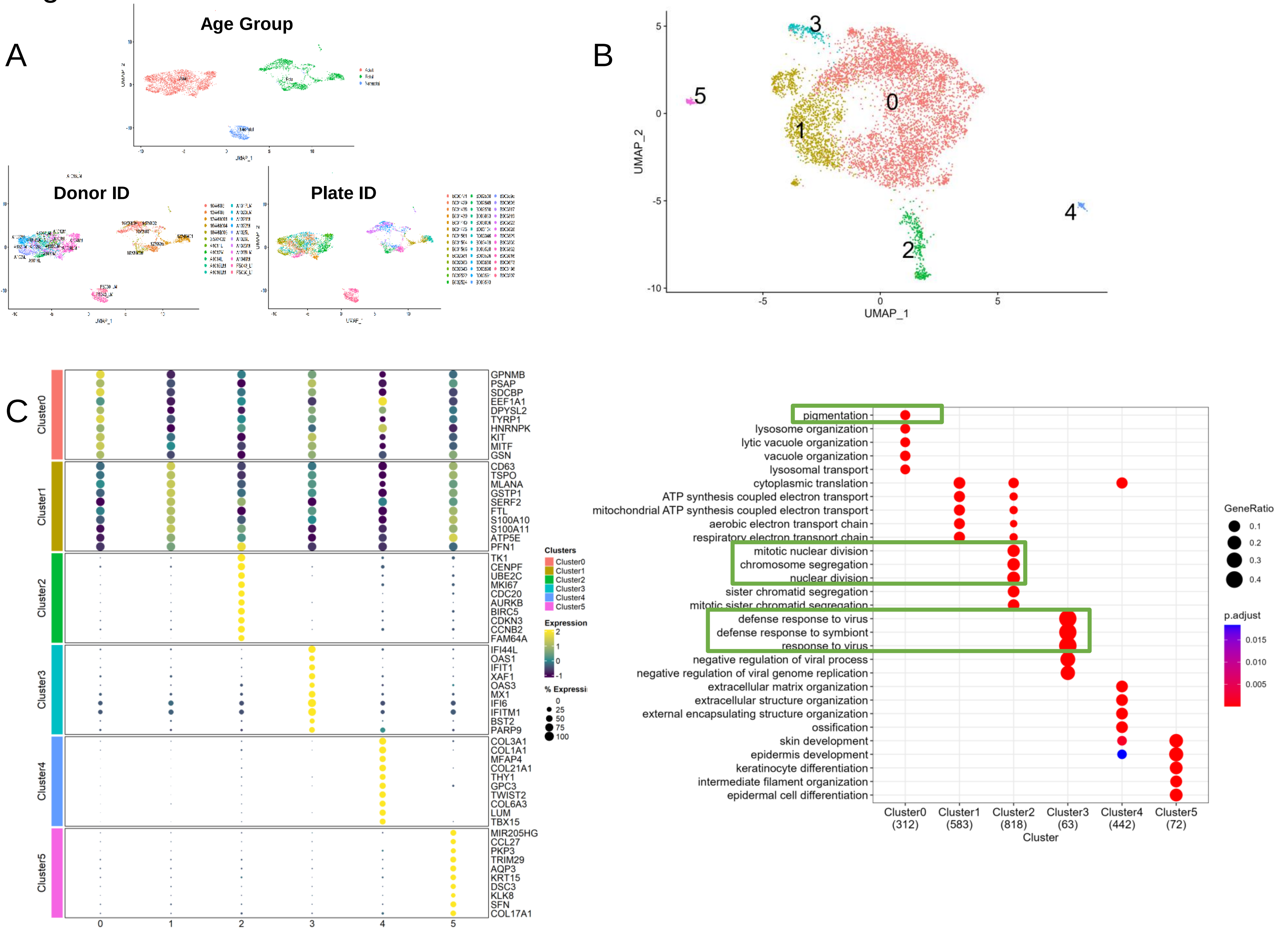

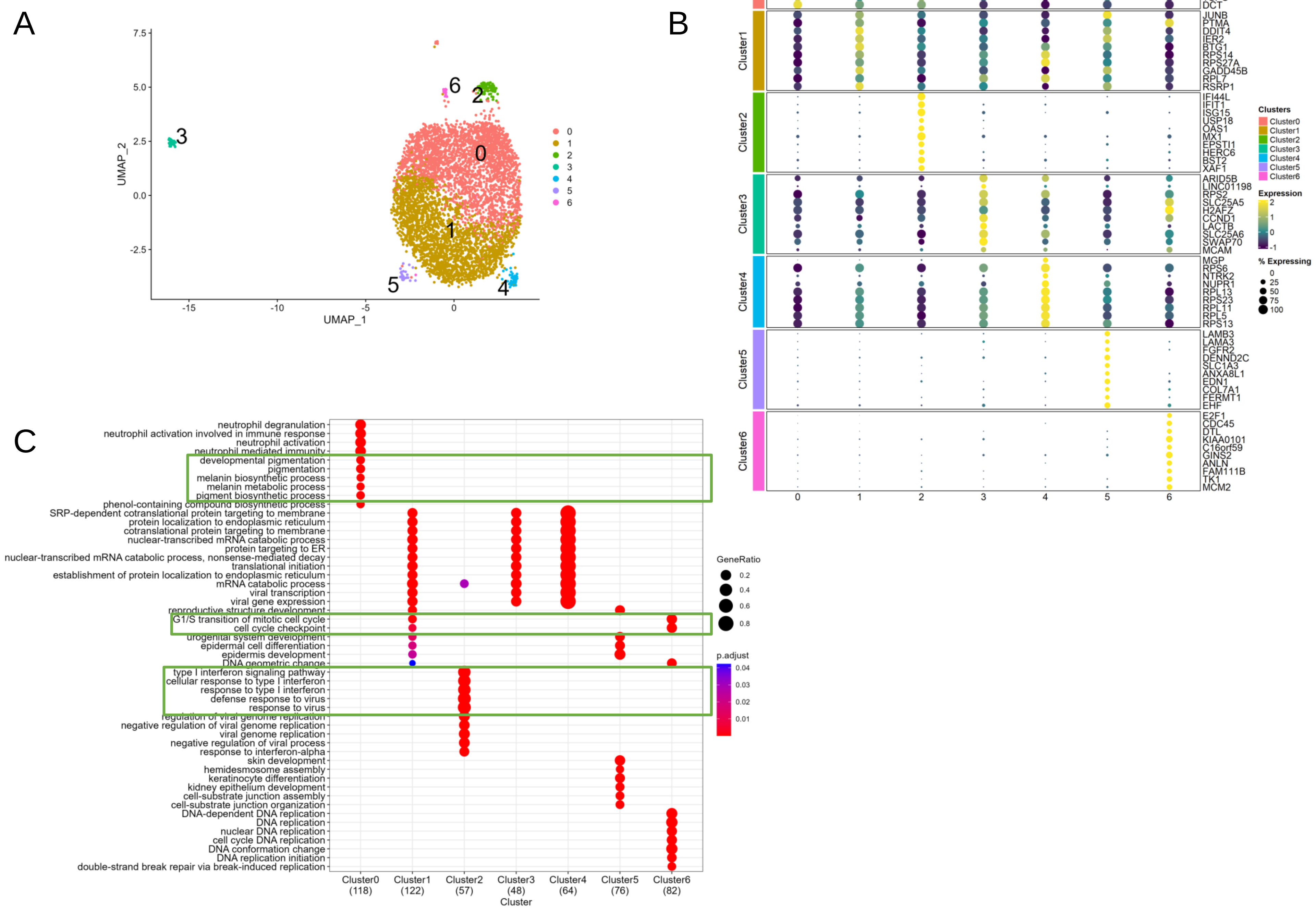

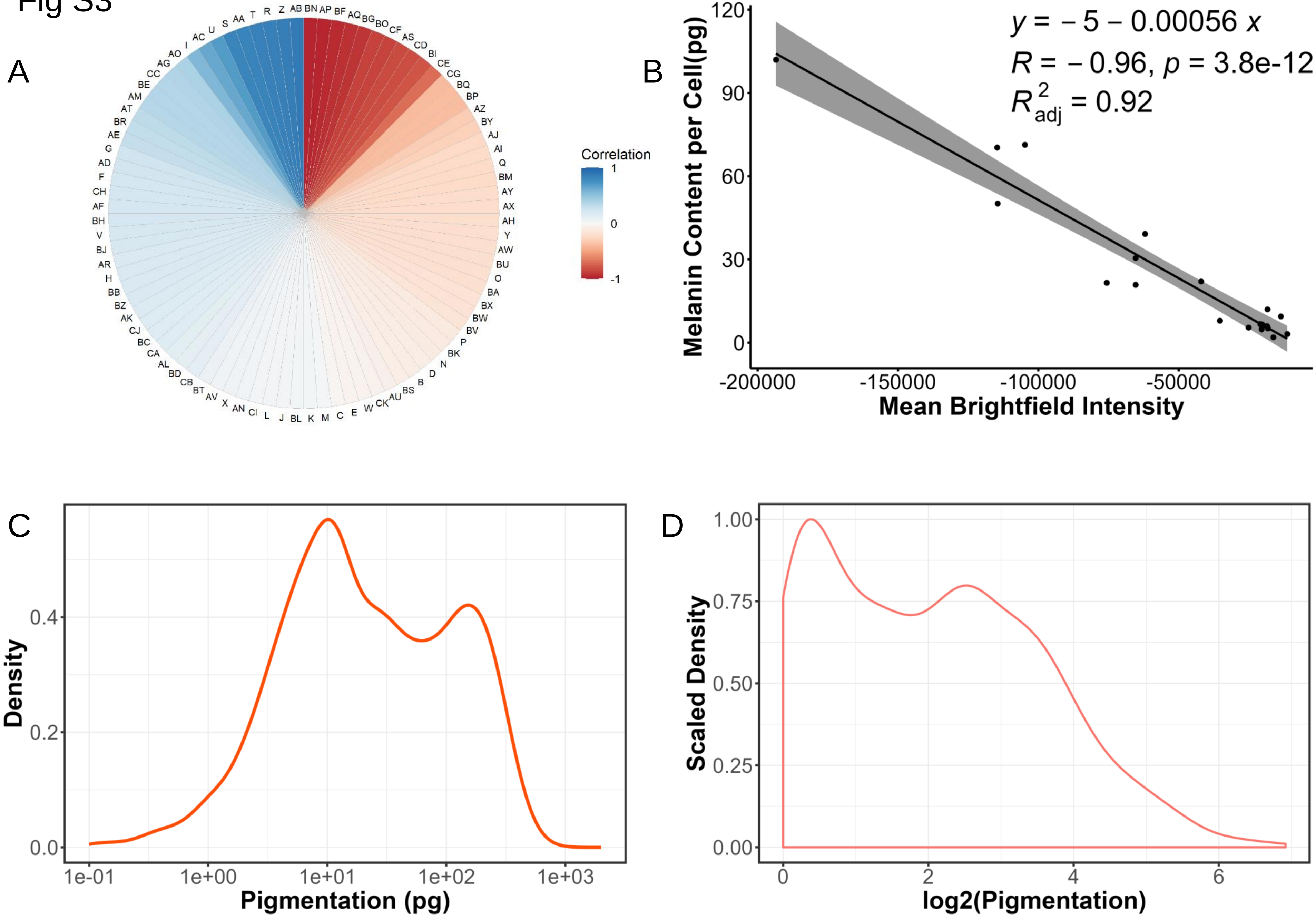

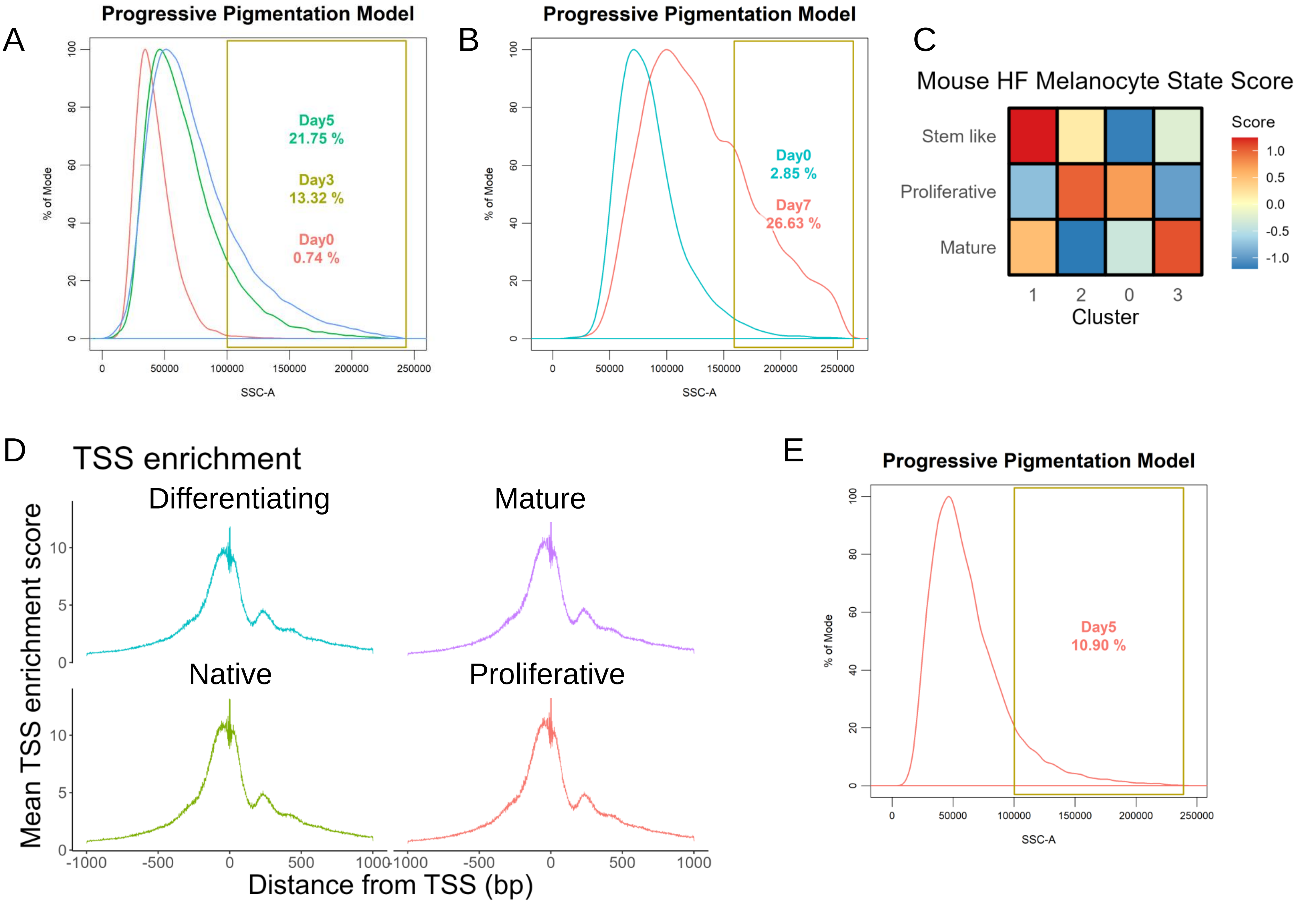

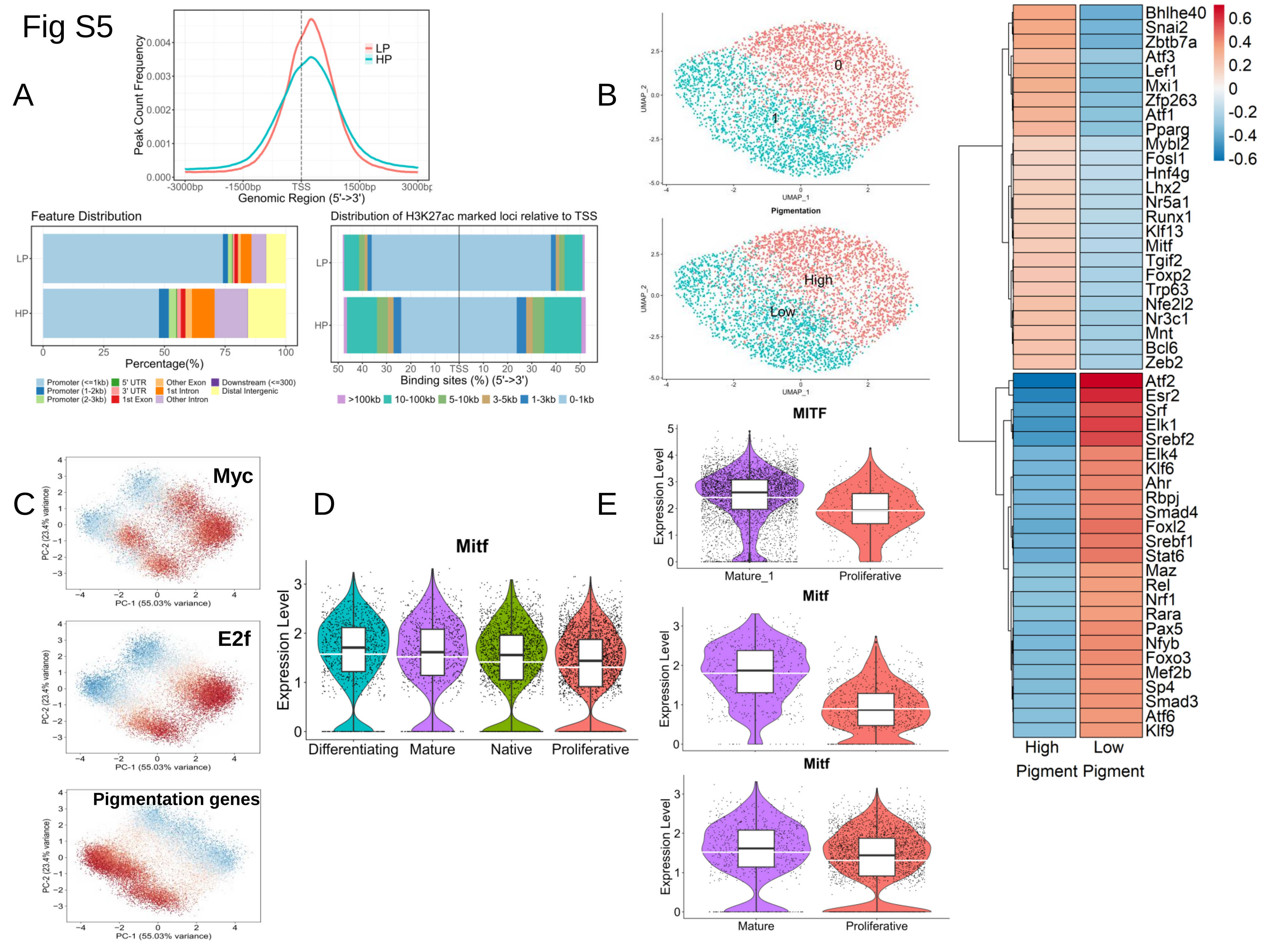
